# Transition of *Staphylococcus aureus* tetracycline resistance plasmid pT181 from independent multicopy replicon to predominantly integrated chromosomal element over 65 years

**DOI:** 10.1101/2025.09.14.675889

**Authors:** Megan A. Phillips, Robert A. Petit, Daniel B. Weissman, Timothy D. Read

## Abstract

Mobile genetic elements (MGEs), including plasmids, phages and genome islands, are major sources of bacterial genetic diversity. The small plasmid pT181 confers tetracycline resistance in bacterial pathogen *Staphylococcus aureus* via an efflux pump, TetK. pT181 was one of the earliest sequenced *S. aureus* plasmids and has been isolated in both clinical and livestock-associated strains for decades, both as an independent replicon and integrated in the chromosome as part of staphylococcal cassette chromosome *mec* (SCC*mec*). Bacterial genome analysis tools and high-quality sequences with metadata are publicly available, but these resources remain underleveraged for examining historical data, especially when studying the spread of MGEs across a species and over time. Using publicly available reads and metadata, we explored the evolution of pT181 over nearly seven decades of samples to identify temporal trends in sequence evolution, copy number changes, and spread across *S. aureus* and beyond. pT181 was prevalent across *S. aureus* (found in 9.5% of 83,366 genomes tested), with a conserved sequence outside of three hypervariable regions. The history of pT181 since 1954 was characterized by spread across strains, significant variation in plasmid copy number of the independent replicon, and increasing frequency of integration of the plasmid into the *S. aureus* chromosome. We identified multiple chromosomal integration locations of the plasmid, including outside of the previously characterized SCC*mec*. We found that pT181 was transferred across *Staphylococcaceae* and into a Gram-negative species. The repeated integration of pT181 into the chromosome may indicate coevolution of the plasmid and the host, potentially to facilitate increased antibiotic resistance.

## Introduction

Plasmids are extrachromosomal DNA molecules capable of autonomous replication that play a major role in horizontal gene transfer (HGT) in many bacterial species. Plasmids have been the most important vectors for the spread of antibiotic resistance genes within bacterial pathogen species since the industrial production of the drugs, starting in the mid-twentieth century ^1,2^. The question of how plasmids themselves evolve, separate from their host, can now be addressed using huge public genome sequence databases and a genomic epidemiology approach. Here, we performed a data-driven exploration of historical evolution and horizontal spread of the small tetracycline-resistance plasmid pT181 in the important human pathogen *Staphylococcus aureus*.

*S. aureus* is a Gram-positive bacterium that asymptomatically colonizes the nares and other body sites of roughly a third of the human population ^3^, but is also able to cause a diverse set of infections ranging from mild to life-threatening. *S. aureus* can be found in both community and hospital settings and is also a significant pathogen of livestock. In a recent paper ^4^, Raghuram et al. (2024) found that an average nucleotide identity (ANI) of 99.5% could be used as a robust threshold to cluster *S. aureus* into 145 distinct strains (numbered consecutively with a “S99.5_” prefix, e.g. S99.5_1, S99.5_2, etc.). This strain threshold is consistent with the gaps in genetic diversity between strains found in other species ^5^. ANI defined strains, defined based on core genome identity, offer more natural boundaries or subspecies groups than traditional allele-based sequence type of clonal complexes such as MLST ^6,7^. In this study, we tracked the transfer of pT181 between the strains defined by Raghuram et al. (2024).

*S. aureus* has undergone waves of evolution of resistance to antibiotics since the introduction of the drugs in the 1940s ^8^, with the most well-studied event being the acquisition of the *mecA* gene on the mobile SCC*mec* elements conferring resistance to second-generation beta-lactams and creating methicillin-resistant *S. aureus* (MRSA) lineages. While much attention is given to beta-lactam resistance, tetracyclines have also been a clinically important treatment option ^2,9^. Tetracycline antibiotics work by binding the ribosome to prevent protein synthesis, arresting growth ^9^. While there is evidence for ancient human use of tetracyclines in Sudan ^10^, the first modern antibiotic in the tetracycline class, aureomycin, was isolated from *Streptomyces aureofaciens* in 1948 ^1^ and entered clinical use soon after. Second-generation tetracyclines were developed in subsequent decades, with doxycycline being introduced in 1967 ^2^. Doxycycline is still widely used in humans for a variety of conditions such as Lyme disease, acne, and STIs, and tetracycline is commonly used in animal agriculture. In recent years, tetracycline-class antibiotics have been used frequently in the doxycycline post-exposure prophylaxis (doxy-PEP) regimen to reduce STIs ^11^, and this use has been linked to increased resistance in off-target bacteria ^12^. Resistance to tetracyclines has emerged through two primary mechanisms: ribosomal protection and efflux pumps^9^. In a similar timeline to MRSA, tetracycline resistance in *S. aureus* emerged in clinical practice soon after the first clinical use of the drug in the 1940s ^13–15^.

Plasmid copy number (average ratio of plasmid replicons to chromosomes in the cell; PCN) is an important aspect of plasmid evolution which can have tradeoffs with horizontal transfer potential and levels of antibiotic resistance ^16^, but remains understudied on a population scale. PCN is regulated by various genetic mechanisms, depending on the plasmid, and can range from a single copy to hundreds per cell ^17,18^. High copy number plasmids can impose a fitness burden on the host due to the metabolic cost of replication and transcription ^19–22^, while low copy number plasmids risk not being inherited by daughter cells without segregation mechanisms encoded on the plasmid ^23^. Larger plasmids often have genes ensuring proper segregation, whereas smaller, high copy number plasmids such as pT181 rely on random diffusion for distribution during cell division ^17,23^. PCN is an evolvable trait of plasmids. For example, elevated PCN of antibiotic resistance plasmids can evolve rapidly following antibiotic exposure ^24^. In addition, PCN is tunable by some bacteria ^25^ and chromosomal mutations can impact PCN ^26^.

pT181 has a circular, 4.4 kbp genome encoding three genes: *repC*, *tetK* and *pre*, and an *oriC* ^27^. It does not encode the molecular machinery to transfer itself into a new recipient lineage and instead must rely on mobilization by a co-resident conjugative plasmid or transduction by phages ^17,28,29^. pT181 confers tetracycline resistance to its host bacterium via the TetK tetracycline efflux pump. Pre facilitates mobilization, but can also play a role in plasmid cointegration ^17,30^. RepC binds and nicks the origin to initiate rolling-circle replication ^31^. pT181 copy number is regulated by countertranscript RNAs binding to the *repC* mRNA to suppress production of RepC, thus downregulating plasmid replication ^32–34^. pT181 was the first *S. aureus* tetracycline-resistance plasmid to be sequenced ^27^ and the plasmid was also significant for its use for multiple seminal studies on DNA replication control ^18,34,35^. Early studies estimated wild-type pT181 PCN to be approximately 22 +/-5 copies/cell ^36,37,34^. It has been known to integrate into SCC*mec* ^38–40^ and is part of some larger staphylococcal plasmids ^41–43^ pT181 has been called by several names in the literature; pT181 is the most common ^44^, but it has also been called pSK52 ^15^, pK34-7-1 ^45^, and rms7 ^46^.

pT181 is an ideal subject for genome-driven evolutionary studies because of its small size and wide distribution. In this study, we screened a collection of over 80,000 *S. aureus* genome sequences for pT181. We developed a novel workflow to use relative read depth from FASTQ to estimate PCN and found evidence that pT181 copy number has declined within *S. aureus* since the first resistant lineages were detected in the mid-twentieth century. This decline was driven in part by integration of pT181 into other genetic structures. We identified repeated pT181 integration into the chromosome, including events outside of the previously characterized region of SCC*mec*.

## Results

### Early pT181 was found exclusively in a single strain, detected later in diverse *S. aureus* genetic backgrounds

Of 83,366 publicly available *S. aureus* Illumina raw data genome projects used in this study ^4^, 7,927 (9.5%) contained pT181, with all isolates having BLASTN ^47^ hit coverage >95% of the plasmid (see Methods) and <5 mutations in coding sequences. There was a reliable date of isolation attached to metadata from 42,522 (51%) of the 83,366 genomes, with the earliest sample from 1884 (SRX2619650) and the latest in our set in 2022. 2,460 isolates contained pT181 and had a valid date, strain assignment, clonal complex (CC) assignment, and SCC*mec* typing data (Fig. S2).

The early pT181 genomes were almost all S99.5_2 ^4^, which are in MLST CC8 ^7^. The earliest dated genome with pT181 was a methicillin-sensitive *S. aureus* (MSSA) isolate collected in 1954. pT181 was exclusively found in S99.5_2 genomes until 1980, when it was detected in S99.5_27 (CC8), integrated into the chromosome as part of type III SCC*mec*. By 2022 (the latest date in our collection), pT181 had been detected in 44 strains. While the number of pT181 genomes collected per year after 1954 increased, overall pT181 prevalence has decreased over time (Fig. 1A) (logistic regression, p < 2e-16). The vast majority of high-quality *S. aureus* samples collected before 1995 were from S99.5_2 (225/242) (Fig. 1B). The subset of high-quality pT181-positive samples collected before 1995 were similarly almost all from S99.5_2 (182/184). Therefore, with this set of genomes, we cannot know if the pT181 *S. aureus* origin was in S99.5_2, or if it was simply detected there first.

**Fig. 1:**
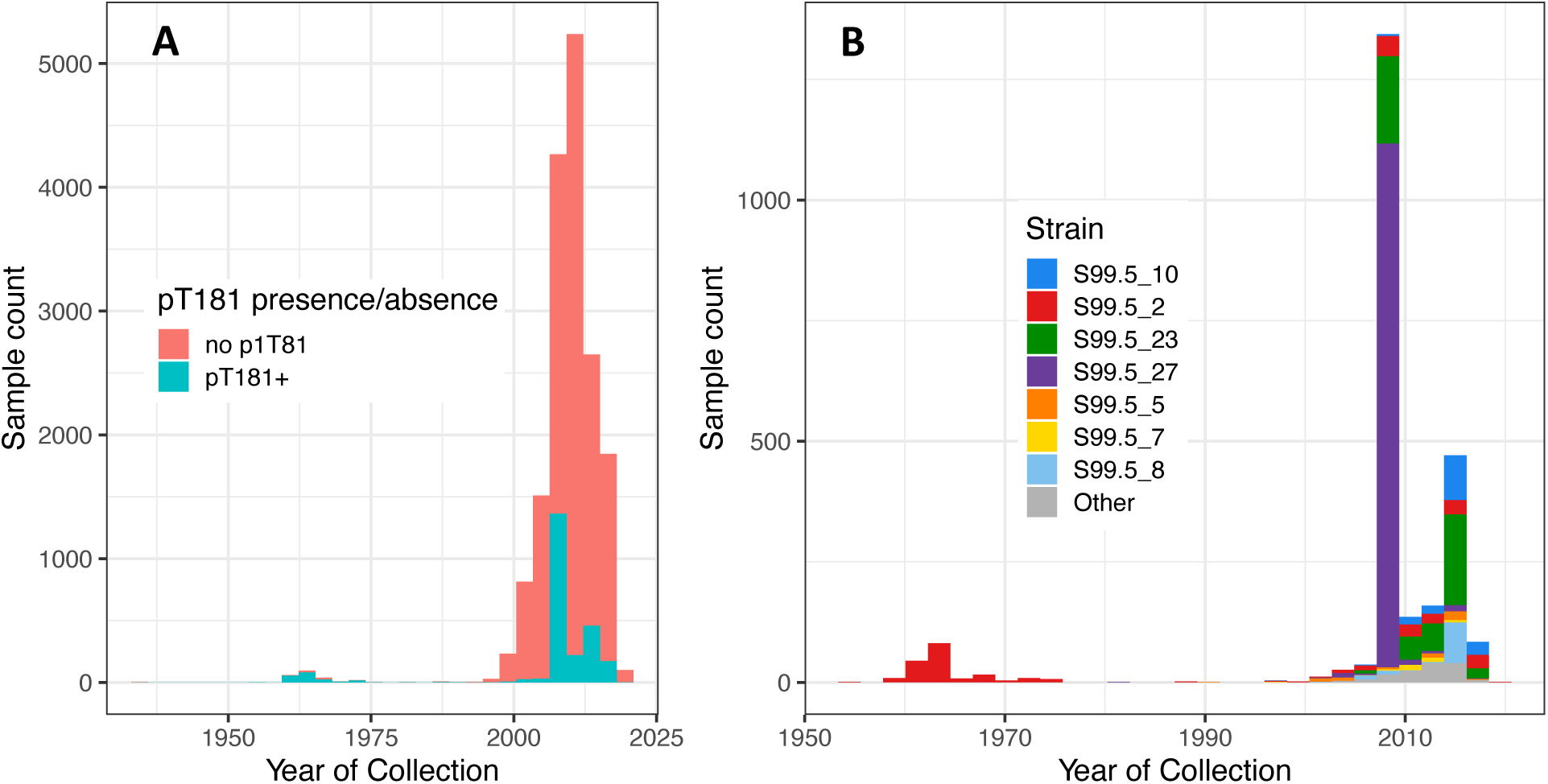
A) Presence/absence of pT181 in high-quality *S. aureus* shotgun genome data over time (n = 16,928). pT181+ isolates comprise less of the sample set over time, though counts of collected samples increase. B) Strain composition of pT181-containing samples has diversified since pT181’s initial detection in S99.5_2.

We also identified pT181 in 61 genome assemblies of 14 other species (File S1). The majority were from *Staphylococcaceae*, but there was also a plasmid found in *Pseudomonas aeruginosa* with plasmid coverage that exceeded the length thresholds for inclusion (see Methods). The *P. aeruginosa* sample was collected from tracheal swabs ^45^, which are frequently taken from cystic fibrosis patients, in whom co-infections with *P. aeruginosa* and *S. aureus* are common. However, HGT between these gram-negative and gram-positive species are rarely documented.

### pT181 sequence variation is concentrated in hotspots outside the coding sequence

Overall, there was a low level of sequence variation in pT181 *S. aureus* and non-*S. aureus* plasmids in this study. 519 sites out of 4,439 sites (12%) on the plasmid had more than one allele in our set of 7,988 sequences (Fig. 2). 664 unique mutations were identified with Snippy ^48^, and 656 (99%) of these are rare, occurring in <5% (<400/7,988) of samples. This variation was concentrated in three regions. The phylogenies of both the whole plasmid (Fig. S1A) and coding sequences (CDS) (Fig. S1B) had a starlike structure with poorly supported branches. The mutation rate in pT181 CDS was slightly lower than most estimates of mutation rates in core chromosomal genes (pT181 = 1.15 SNPs/Mbp/year, SE = 0.3; chromosome = 1.2 - 4.4 SNPs/Mbp/year^49–53)^ when calculated via linear regression of SNP accumulation over time. While the relationship between time and mutation accumulation was significant, time since sample collection was a poor predictor of CDS SNP count (linear regression, p = 0.0001324, adjusted R^2^= 0.006) (Fig. 3). As a result, we are unable to approximate dates for undated samples based on plasmid sequences. This regression excluded the intergenic regions and insertion mutations in *repC*, which had higher levels of variation. There was no evidence of recombination with a pairwise homoplasy index (PHI) test ^54^ of all (n = 7,988) aligned plasmid sequences (p = 0.39), as expected given that the low level of sequence diversity greatly reduced the power of the test. We concluded that pT181 entered the *S. aureus* population since the divergence of the strains ^53^, given that most pT181 plasmids have identical CDS (Fig. 3).

**Fig. 2:**
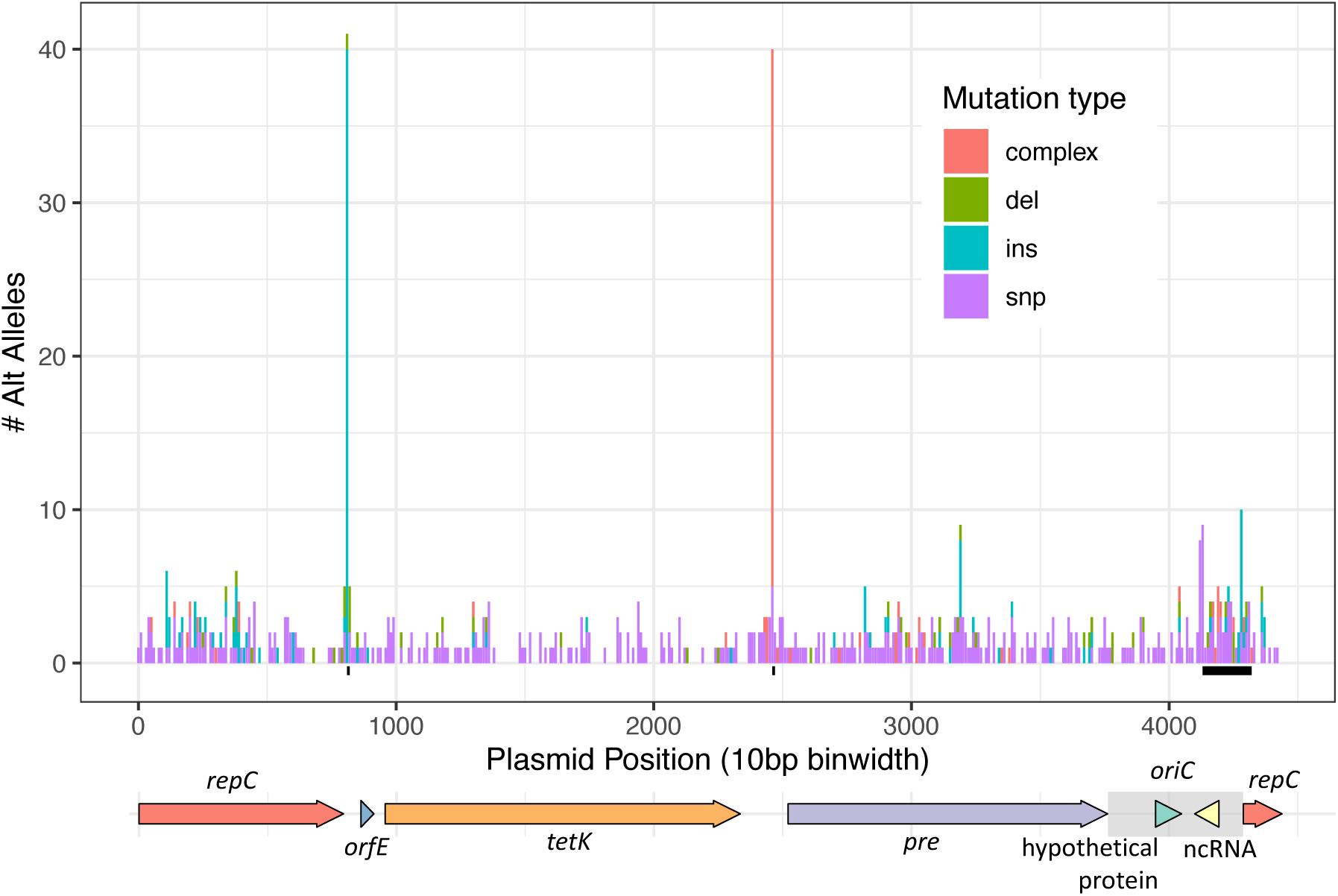
pT181 is characterized by hotspots of sequence variation. Hypervariable regions have a large number of alleles present. Each hypervariable region is characterized by a set of mutation types and is marked by a black dash underneath the x-axis. Binwidth = 10 bp.

**Figure 3:**
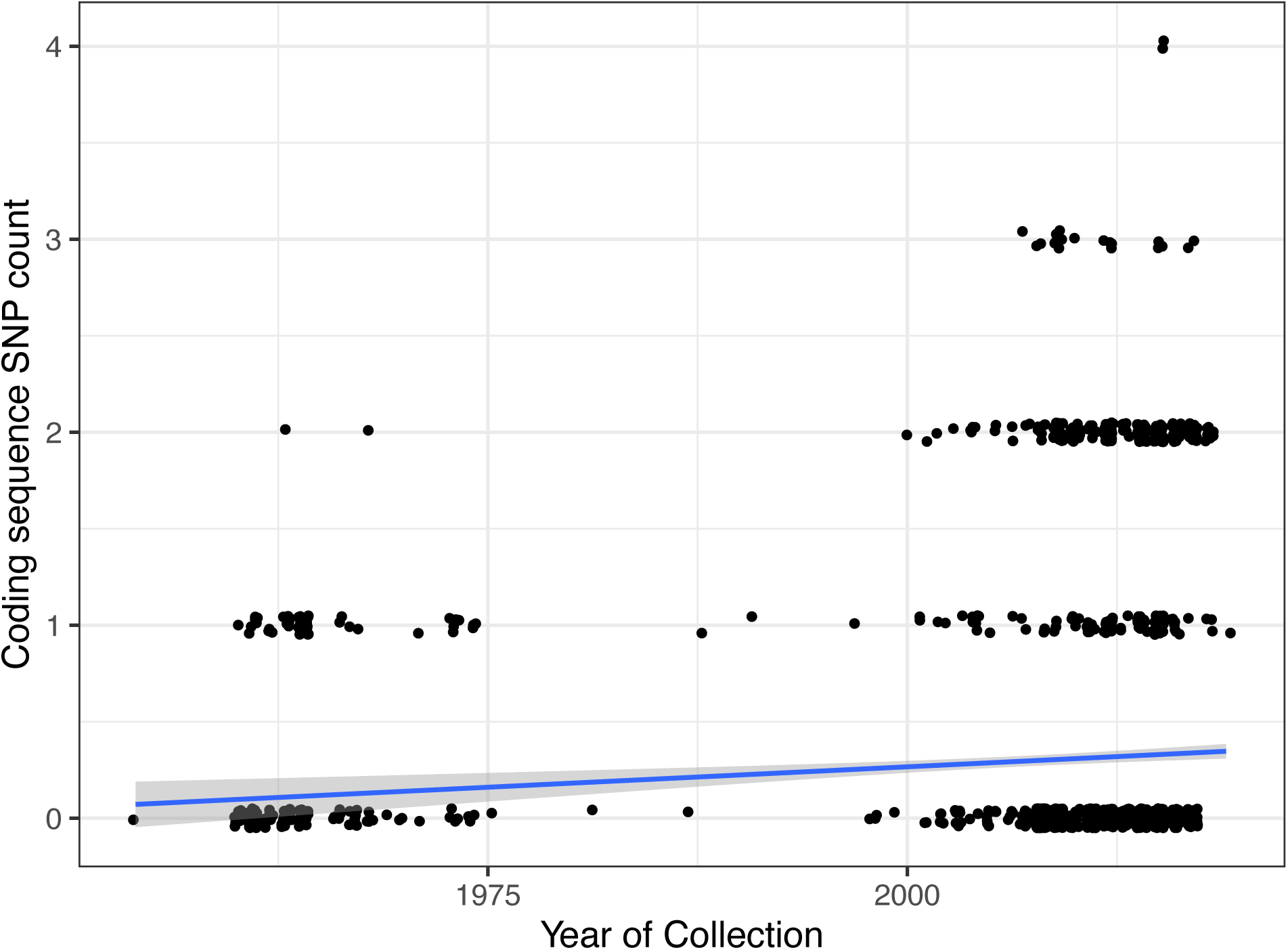
pT181 CDS SNP accumulation over time relative to the 1954 isolate, calculated from Snippy output.

While the majority of the pT181 sequence was highly conserved, three regions of increased variability were present on the plasmid: 1) the intergenic region between *repC* and *tetK*, (coordinates 814 to 817 on accession DRX100326), 2) the intergenic region between *tetK* and *pre*, and (coordinates 2460 to 2466 on accession DRX100326) 3) in *oriC* and early in *repC* (coordinates 4133 to 4313 on accession DRX100326). Mutations immediately before, and at the 5’ end of *tetK* are found exclusively in single-copy isolates (see later section on PCN). The spikes coincide with known plasmid cointegration sites ^42,43,55^.

pT181 in *S. aureus* was highly similar to pT181 found in other species. The *S. aureus* and non-*S. aureus* plasmids were all in a highly similar cluster (5 or fewer CDS mutations relative to the 1954 isolate), with the exception of *Staphylococcus pseudintermedius* (CP128253.1), which had 12 CDS mutations (8 SNPs, 4 complex mutations). Only 5 non-*S. aureus* isolates contained a mutation exclusive to non-*S. aureus* isolates, but these were at low prevalence, with a SNP present in 2 isolates and a complex mutation present in 3 isolates. These results indicate that the non-*S. aureus* pT181 sequences did not constitute an outgroup to the *S. aureus* pT181 sequences.

### pT181 is most frequently found at a single copy per cell

We noted a skewed distribution of PCN in gold-quality *S. aureus* pT181 isolates, with a wide range of PCN estimates (Fig. 4A) and a strong peak of pT181 PCN = 1 (Fig. 4B), consistent with chromosomal copy number.

**Fig. 4:**
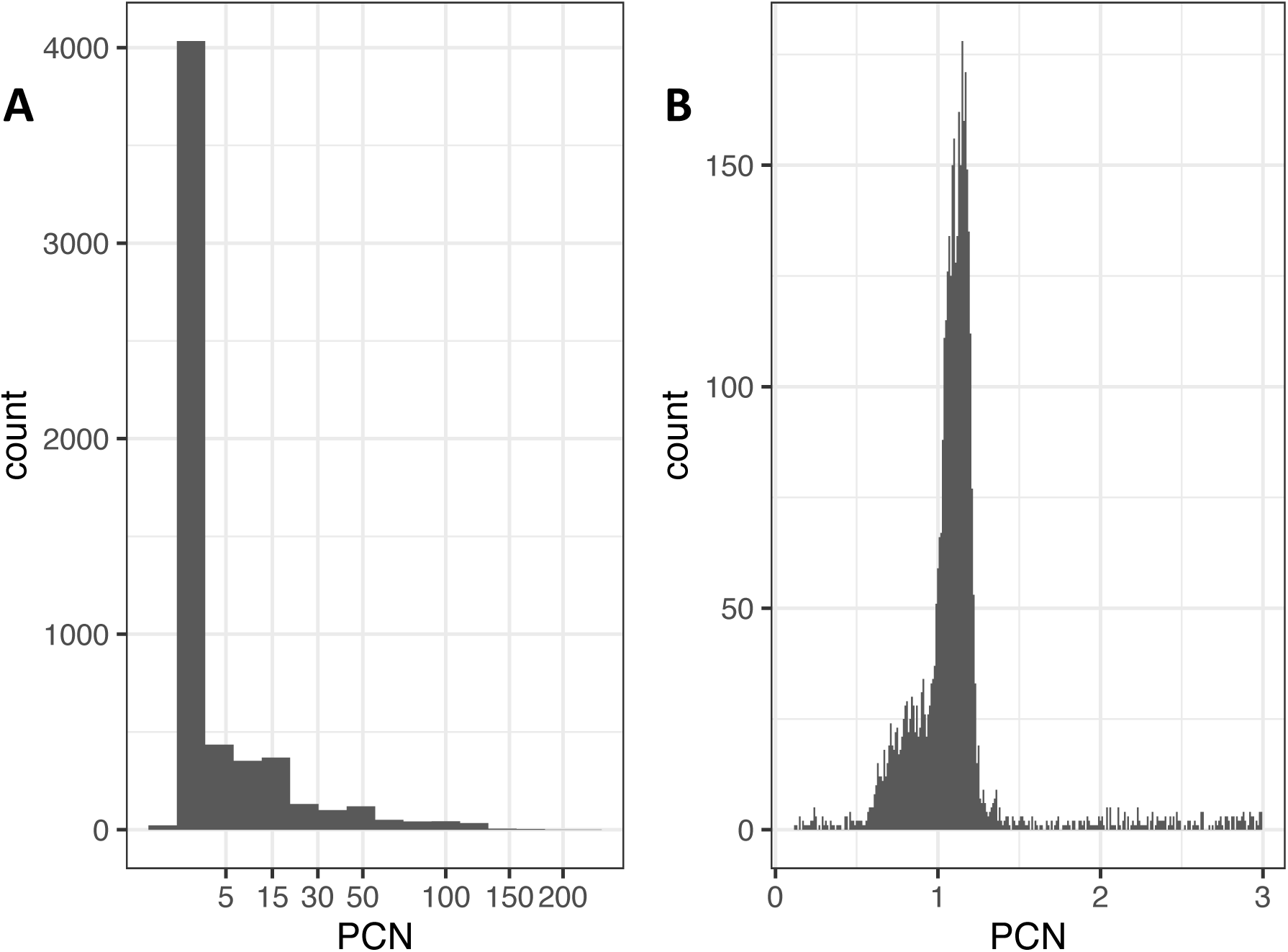
A) Overall PCN distribution for gold-ranked, pT181+ isolates. The majority of pT181 isolates in our dataset are single-copy. Given that pT181 lacks segregation mechanisms, this suggests that the plasmid has become integrated into the chromosome. B) Subset of PCN distribution, enlarged for 0-3 copies/cell.

pT181 can be present on the chromosome as part of type III SCC*mec*, which confers both methicillin and tetracycline resistance ^38,39^. We found that the average PCN of type III SCC*mec* strains was 1.16 with a range of 0.71 to 1.43 (see Methods), matching the expected distribution given SCC*mec*’s position near the origin. Based on this result, we designated any sample with pT181 PCN < 2 as single-copy. Of 2,460 genomes with valid date, strain assignment, CC assignment, and SCC*mec* typing data, 1,863 (76%) were single-copy (PCN 0.147 - 1.97; mean = 1.06, sd = 0.186). We suspect that these single-copy pT181 were integrated into the chromosome or a low copy number plasmid. Instances of integration into the chromosome in other types of SCC*mec* have been previously documented ^40,55,56^. We found that 1,131/1,863 single-copy pT181 (61%) were present in SCC*mec* type III isolates, 695/1,863 (37%) were in other MRSA strains, and 37/1,863 (2%) were in MSSA, i.e. lacking a SCC*mec* cassette.

### pT181 has inserted into multiple chromosome sites

There were 19 isolates with pT181 present on the same contig as a known chromosomal gene in the SRA-assembled genomes, comprising 6 unique insertion sites as defined by directly flanking genes. All had pT181 present in a single copy number. The isolates were distributed across multiple strains: 5 S99.5_5 (CC5), 2 S99.5_2 (CC8), 8 S99.5_23 (CC398), 1 S99.5_10 (CC1), and 3 with unassigned strain (CC1 & unassigned). One isolate was MSSA, 3 were SSC*mec* type IV, and 15 were SCC*mec* type V. To our knowledge, this is the first documented example of chromosomal pT181 in MSSA. The integrated pT181 were flanked by insertion sequences (IS): IS*6*, IS*Sau6*, or IS*257* transposases, or by one transposase and a contig break. Integrated pT181 was present in multiple configurations and positions in a genome, and there were no nearby chromosomal genes common to all integration sites, other than some form of flanking transposase (Fig. 5). Some pT181 integrated with *repC* positioned nearest the chromosomal flanking genes, and other isolates *tetK* nearest, due to variation in the site of plasmid cointegration. Some *S. aureus* strains had multiple different insertion sites, indicating repeated integration following introduction into a strain. There were plasmids with identical sequences that were both single and multicopy, lending further support to our finding that the integrations have happened repeatedly over the course of pT181 history.

In 337 *S. aureus* complete genomes ^4^, we found that 14 (4.2%) contained pT181. 7 of these were integrated into the chromosome, 4 were integrated into larger plasmids (total plasmid length: 21-44kbp), and 3 were free in the cytoplasm. The integration of pT181 into larger plasmids may result in low pT181 PCN, as pT181 could take on the copy number of the larger element. Ultimately, if the copy number controlling sequence of one plasmids is not disrupted during the integration process, the PCN of hybrid plasmids may be determined by replicon dominance ^57^. The chromosomally integrated copies of pT181 occurred in positions 55-88kbp in these genomes, near SCC*mec*, with both positive and negative sense orientations. The region in which pT181 integrated into the chromosome in these genomes is populated primarily by rare and strain-diffuse genes ^4^ (Fig. S3). The presence of pT181 in larger plasmids, which tend to be lower copy number ^58^, as well as integration into the chromosome, indicate additional routes to evolve low copy number beyond regulation of free plasmid.

**Fig. 5:**
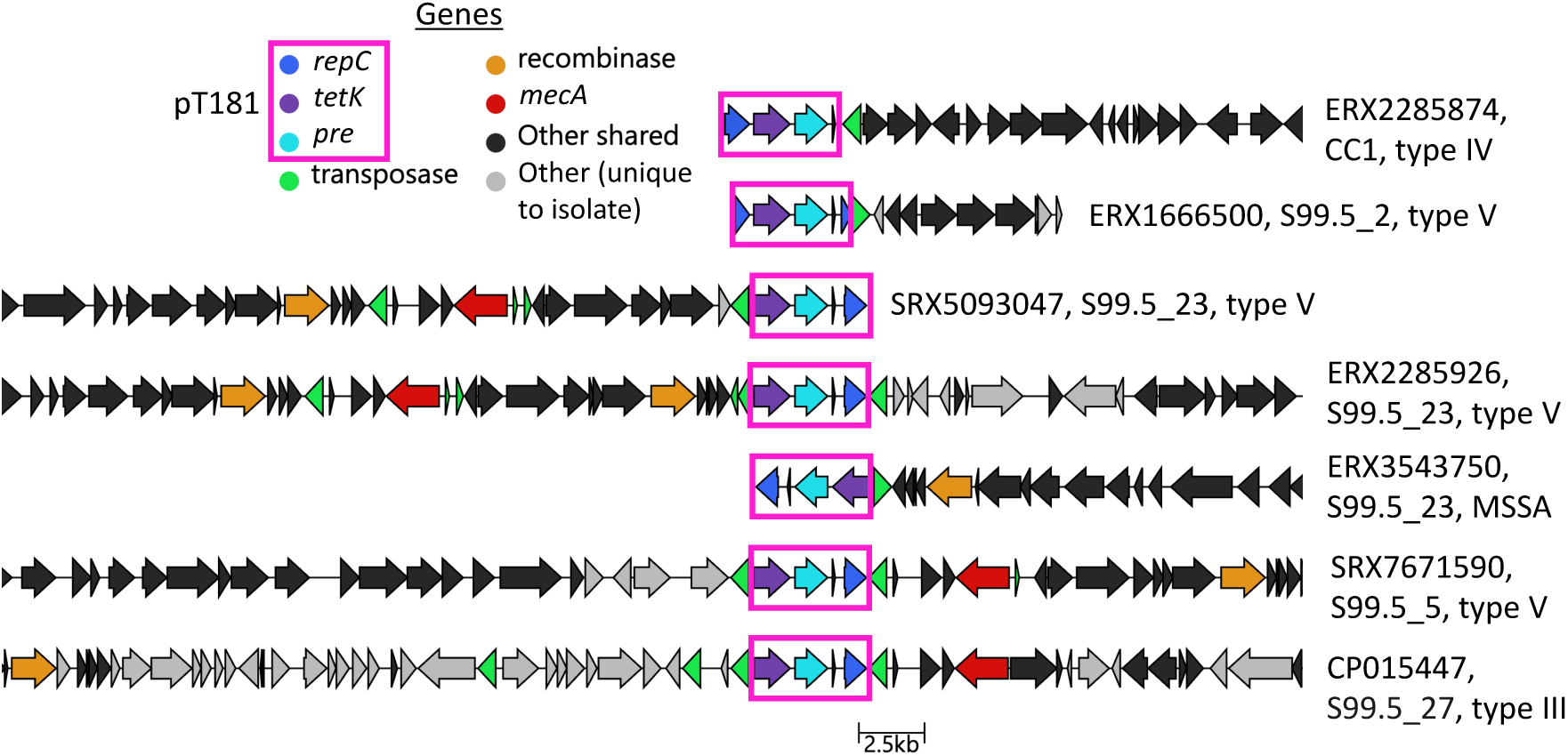
There were 6 non-type III SCC*mec* configurations of accessory genes directly surrounding the integration site of pT181 contigs across multiple strains in the SRA isolates. This shows that the plasmid has integrated repeatedly into the chromosome, given that *S. aureus* is highly syntenic and accessory genes tend to cluster in specific regions of the genome ^4,59,60^.

### pT181 PCN has declined over time

pT181 PCN declined over time from the 1950s through the end of collection of our sample set in 2022. This is the result of two trends: (1) increase in single-copy isolates over time and (2) lower PCN over time in multicopy isolates. We performed separate regressions to assess prevalence of single-copy versus multicopy pT181 over time and change in copy number in multicopy isolates over time.

Over the period for which we have dated samples, pT181 was increasingly found in single-copy form (Fig. 6A) (logistic regression, p < 2 x 10^-16^). Pre-1995 samples were mostly multicopy (8.2% single-copy), while 81% of samples from 1995 or later were single-copy. Single-copy pT181 was not evenly distributed across the strains (p < 2 x 10^-16^, Chi-squared test statistic = 2052), with some strains being mostly or entirely single-copy (Fig 6B). S99.5_27, all isolates of which are type III SCC*mec* in our dataset, constitutes a substantial portion of single copy isolates (1,131/1,863), and is responsible for the collection peak in 2008. However, even with this strain excluded from analysis, the majority of the remaining pT181 isolates are still single-copy (55%; 732/1,329) and the increase in single-copy isolates over time is still significant (logistic regression, p < 2 x 10^-16^).

**Fig. 6:**
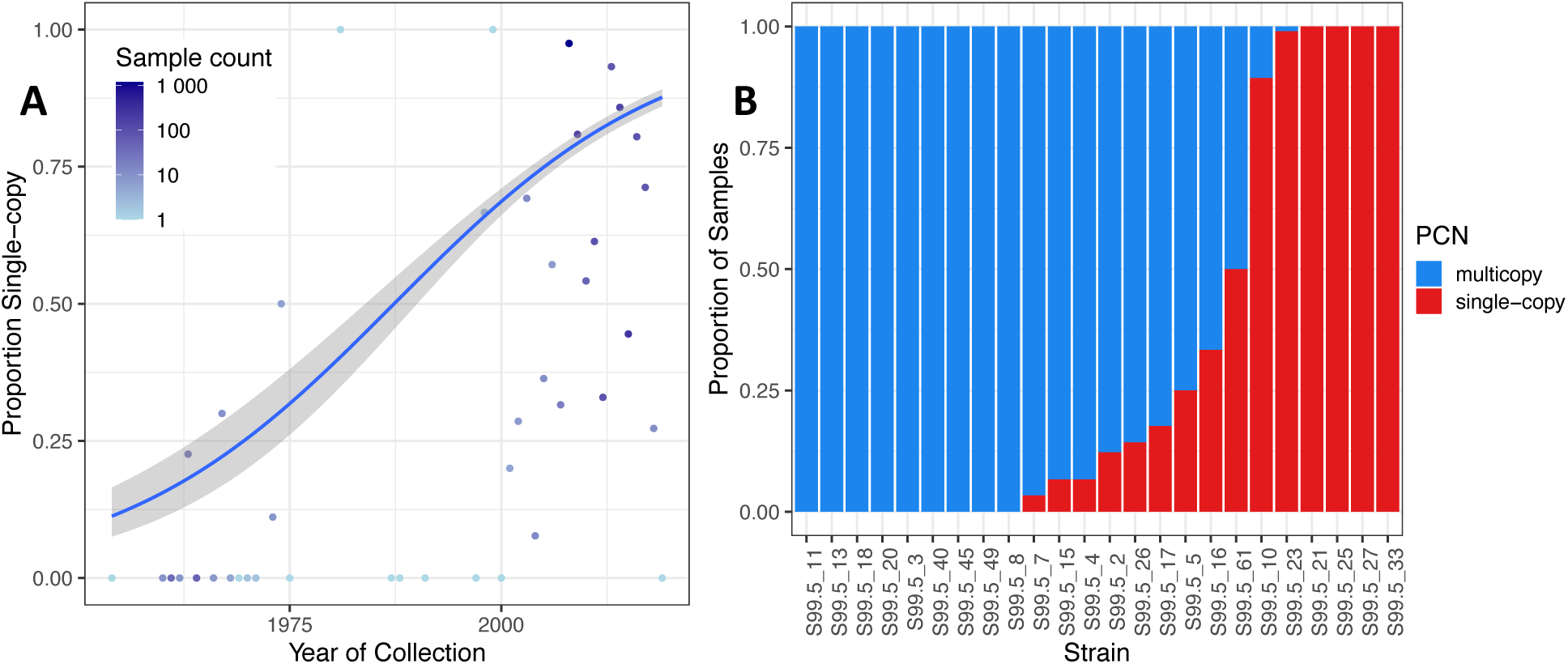
Proportion of pT181 isolates present at single-copy number has increased since initial detection and varies across strains. A) Proportion of single-copy pT181 increased over time. B) Single vs. multicopy pT181 across strains, for strains containing >= 2 pT181+ isolates in (n = 2,460) set. Strains are generally dominated by either single or multicopy pT181.

Copy number of cytosolic plasmid could be influenced by time or strain. We used a Gamma log-link generalized linear mixed effects model (GLMM) to assess the impact of date of collection and strain background on PCN in multicopy isolates, while controlling for the impact of technical variation using BioProject (see Methods). There has been a decline of PCN over time in these multicopy isolates (p = 3.30 x 10^-12^). No strains were significantly different from our reference, S99.5_8 (mean PCN = 17.4, SD = 4.35) (treatment vs. control test, emmeans package, p > 0.05, BH multitest correction). The effect of time is still significant if S99.5_2, the strain with the oldest samples and highest PCN values, is excluded from the analysis (p = 0.035). We also assessed the potential impact of DNA extraction techniques on PCN estimation and found that they did not change our conclusions (see Methods). The PCN decline over time within S99.5_2 remains significant when assessing only within samples with identical extraction method, sequencing center, and sequencing instrument (linear regression, p = 5.9 x 10^-16^). The decline in multicopy PCN over time indicates possible evolution of the plasmid, chromosome, or both, toward lower PCN.

**Fig. 7:**
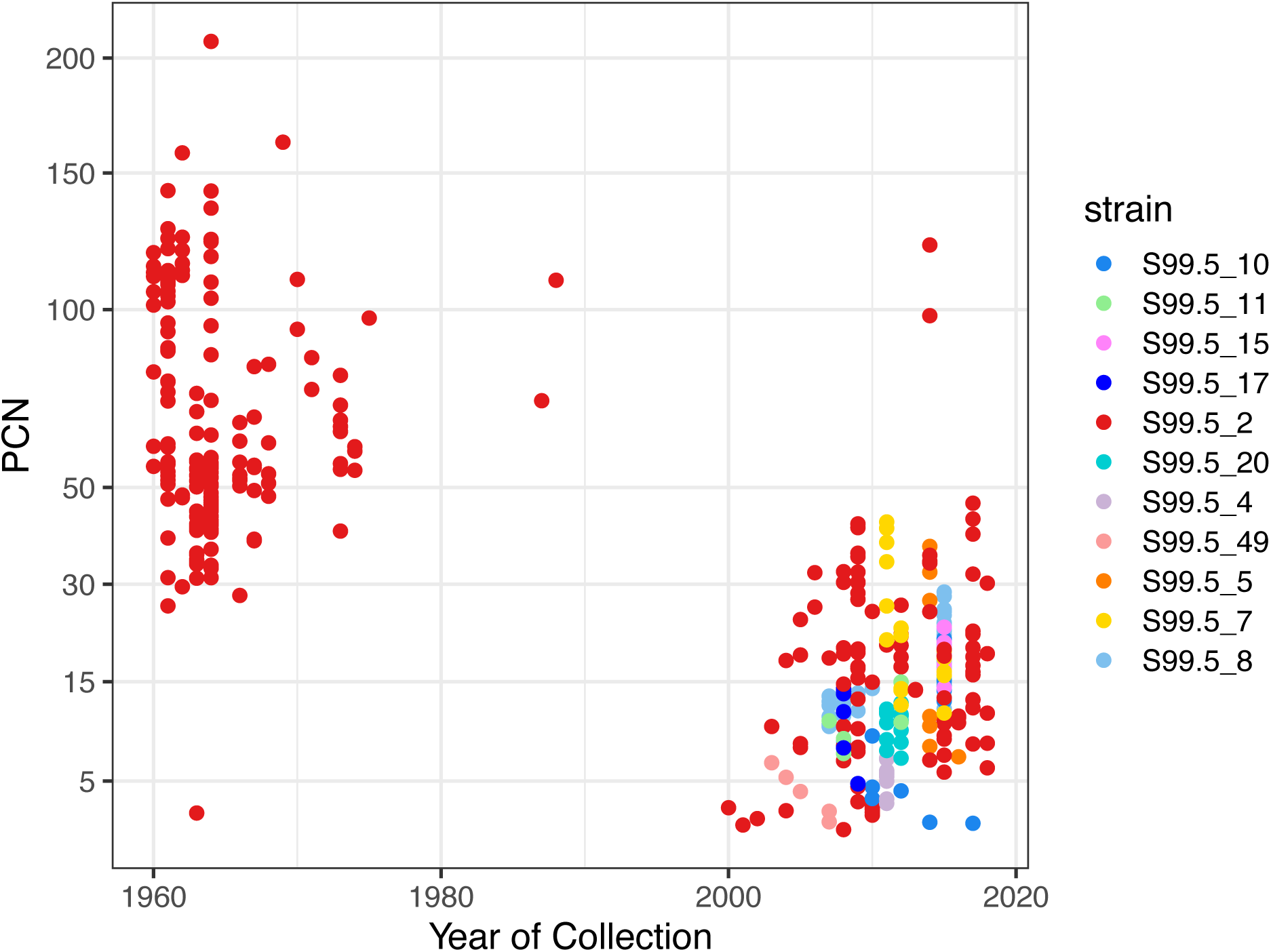
PCN of multicopy isolates has declined over time. Strains’ PCN are not significantly different from reference strain, S99.5_8 (GLMM, treatment vs. control posthoc, BH multitest correction). The effect of time is significant even with S99.5_2 excluded.

### Identical plasmids can have variable plasmid copy numbers

pT181 copy number control mechanisms are located on the plasmid, which regulates its own copy number ^34^. However, previous research has identified chromosomal variants impacting pT181 PCN ^26^. In our set of isolates, we find that identical pT181 isolates can have a wide range of copy numbers (Fig. 8). This suggests that there are factors other than copy number control mechanisms on the plasmid, such as host genetic background or environment, influencing pT181 copy number in our dataset.

**Fig. 8:**
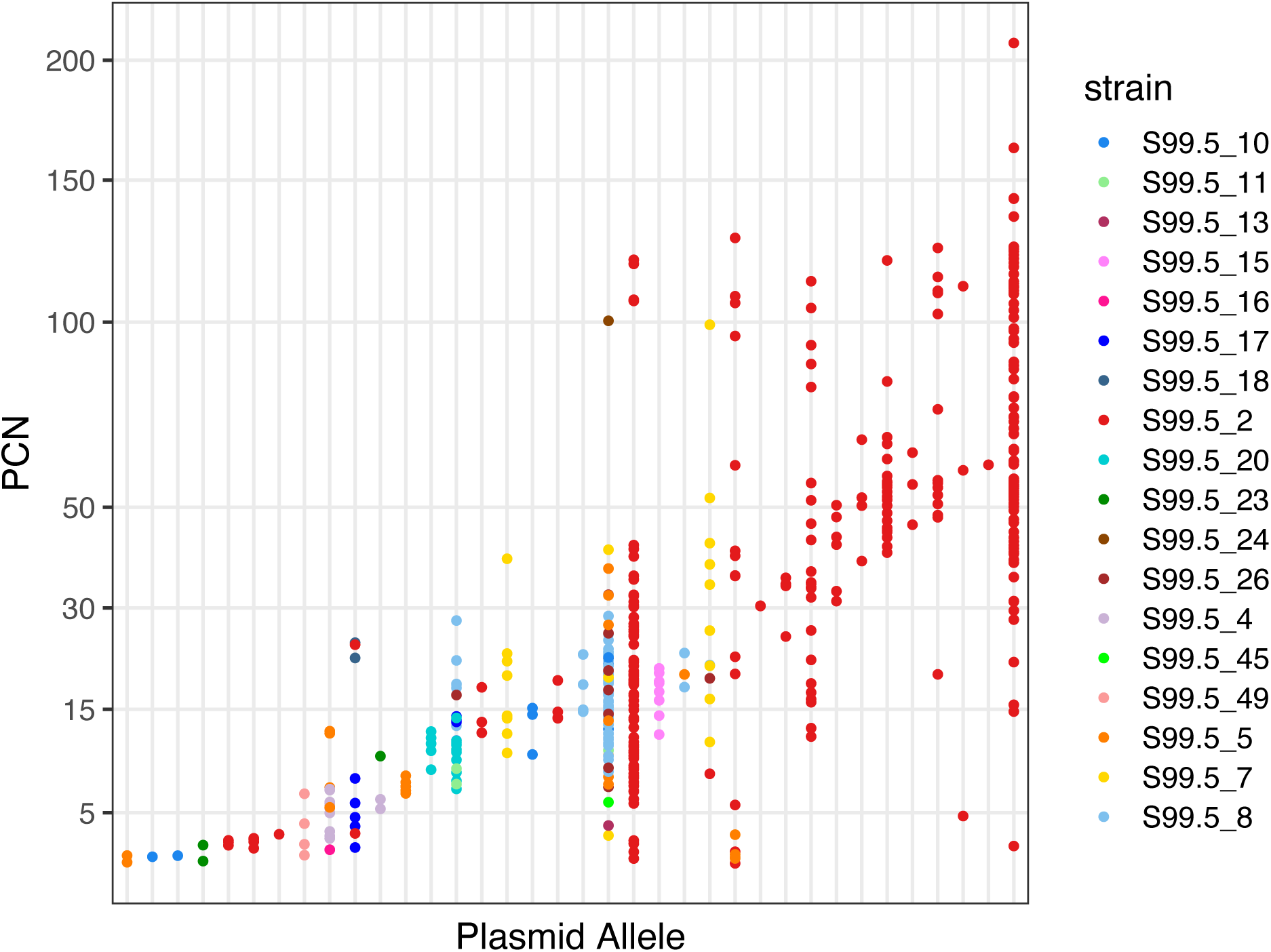
Multicopy PCN varies within identical plasmid sequences and strains, indicating that there are environmental or genetic effects occurring at the substrain level which impact pT181 PCN. Plot shows all unique plasmid sequences (across all sites) occurring at least twice as multicopy isolates in the (n = 2,460) dataset.

## Discussion

This is the first study to use the thousands of available public genome sequences to look at variation of PCN of a single plasmid. We estimated the PCN of multicopy pT181 in high-quality isolates to be 2 to 208 (n = 2,460) with a median of 19 copies/cell (Q1 = 11 copies/cell, Q3 = 47 copies/cell) (Fig. 4A). Our median PCN is consistent with experimental studies for this plasmid’s replicon using qPCR ^61^, relative resistance to antibiotics ^18^, and fluorescence densitometry ^62^. Relative read depth has been previously used for ploidy estimations in cancer and yeast ^63,64^, but we have shown here that it can be accurately employed to estimate copy number of MGEs or other contigs in bacterial genomes, given sufficient knowledge of the genome such that a set of gene or genes can be used as the known single copy regions to compare the unknown region(s). The genomic-based PCN method can be easily applied to thousands of already-collected samples, without the need for specialized qPCR assay development.

In addition, these methods are not culture dependent and could be deployed in metagenomic samples given sufficient quantity of the host in a metagenome sample. We do not have control of growth, extraction, or other preparation conditions that might influence plasmid copy, so we cannot account for all factors that may contribute to the measured copy number. However, in our analysis, we do not find any impact of extraction techniques on PCN. In addition, *S. aureus* is usually grown overnight to stationary phase in rich media before genomic DNA extraction, and we therefore expect reasonable consistency in technique for many protocol steps.

We observed a wide range of pT181 multicopy number PCN’s, outside of the canonical values of approximately 22 copies/cell ^36,37,34^. Cointegration with other plasmids could contribute to variation in pT181 PCN ^57^. Previous investigation into the control of pT181 copy number has discovered the mechanisms present on the plasmid itself ^34^. PCN is known to be controlled by pT181 antisense ncRNA binding to the *repC* transcript, which modulates RepC production and therefore plasmid replication ^32,37,65^ when the plasmid is free in the cytoplasm. Variation that affects plasmid *repC* expression and/or RepC binding to *oriC* can impact the average number of copies per cell ^18,35,41,65^. Previous research has identified chromosomal mutations capable of elevating pT181 copy number ^26^ by reducing countertranscript synthesis and increasing RepC production ^66^. Chromosomal loci could may also influence pT181 PCN through mechanisms such as RNA degradation, but the strain-level and below-strain-level variation of the PCN phenotype has not been an area of intense research, and it is likely that there are additional genes involved in plasmid regulation. At a population level, we find that the strains vary in their average copy number, but this difference was not statistically significant following post hoc testing (GLMM, BH post-hoc, all p-values > 0.05). We also find that identical plasmids can have different multicopy PCN, even within the same strain, emphasizing the need to further investigate genetic contributors on the chromosome and interactions between the chromosome and plasmid. Further research on the genetic basis of variation of pT181 PCN and variation in relative levels of TetK production will help explain how this plasmid-bacterial symbiosis has adapted to the challenge of large scale antibiotic usage.

pT181 PCN declined over time, driven in part by the increased frequency of chromosomally integrated plasmid. This is consistent with historical reports of early detection of multicopy pT181, followed by the tetracycline resistance phenotype becoming chromosomally encoded in subsequent decades ^67^. While it was previously known that pT181 could integrate into the chromosome as part of SCC*mec* elements, it is a new finding from this study that the plasmid is at approximately chromosomal copy number (i.e. single-copy) in the majority (76%) of *S. aureus* isolates. The majority of single-copy pT181 is found in SCC*mec* type III isolates, with pT181 integrated into SCC*mec*; however, even with these excluded, more isolates are single-copy than multicopy. We also identify integration into other SCC*mec* types, as previously described in the literature ^40^. Integration into the chromosome in MSSA isolates is previously uncharacterized in the literature to our knowledge. Integrated pT181 in our dataset are flanked by insertion sequences (IS), or by an IS and a contig break. IS elements have been previously documented to insert in regions of pT181 that we have characterized as hypervariable^42,43^, and this may account in part for the spikes of mutations occurring in these regions in our analysis. IS*6* and IS*6*-like transposases, which flank integrated pT181 in our data set, have been known to integrate multicopy plasmids into the chromosome; IS*257* has been previously found to integrate pT181 into larger plasmids and into the chromosome as part of SCC*mec* ^38,39,42,68–72^.

The decline in PCN over time may be due to selection to reduce the fitness burden on *S. aureus* cells that comes from replicating larger amounts of plasmid DNA. An analogous argument for lower fitness penalty driving the transition of *S. aureus* from larger SCC*mec* type I to smaller type IV has been put forward ^8^. Adaptation could arise through IS-mediated integration into the chromosome or large plasmids, and/or mutations on the plasmid or chromosome replication control circuits to lower PCN. In some scenarios, high PCN may be advantageous as cells with higher PCN of pT181 have higher tetracycline minimum inhibitory concentration (MIC) ^69^ and previous studies have identified gene dosing as a mechanism to increase MIC ^16,24,73,74^. The high PCN in early S99.5_2 isolates may have been permitted because of the benefit of tetracycline resistance, and the decline over time may have been adaptive for long-term maintenance. pT181’s decline in PCN brings it from median multicopy PCN of 55.7 copies/cell pre-1995 to 14.5 copies/cell post-1995 (n = 2,460), a decline from roughly 9% to approximately 2% of the chromosome’s size. The post-1995 median is in line with recent research on plasmid copy number across taxa, which found that plasmids occupy roughly 2.5% of the chromosome’s size ^57^. There may also be genetic adaptations maintaining high levels of *tetK* transcription despite lower PCN. Simpson et al. (2000) ^69^ identified the appropriation of an IS*257*-derived hybrid promoter, which had higher activity than the native pT181 promoter, rendering the integrated copies more tetracycline resistant than the multicopy isolates. In this way, bacteria with integrated pT181 could have been selected for both because of their increased tetracycline resistance due to the hybrid promoter and reduced the cost of maintaining a multicopy plasmid. High levels of *tetK* transcript have been shown to be toxic in *E. coli* ^75^, so there may be a limit to benefits from high levels of transcription with respect to both resources used and impacts on the cell. An alternate but not mutually-exclusive hypothesis is that it was advantageous for *S. aureus* to integrate the plasmid to allow co-residence of another advantageous, but incompatible cytoplasmic plasmid. Ultimately, we are unable to conclude for certain why the pT181 PCN has shown such a marked decline since its detection in mid-twentieth century.

In our study, pT181 was initially detected in S99.5_2, with appearance in other genetic backgrounds occurring decades later. Extensive spread across *S. aureus* has occurred, as well as spillover into other species. The first recorded detection of tetracycline resistance via pT181 dates to 1953 ^15^, very soon after the modern discovery of tetracyclines ^1^. It is likely that antibiotic use was the major selective pressure increasing the incidence of pT181+ isolates since the 1950s. Our finding that pT181 CDS were almost identical across the collection period suggests recent expansion from a single introduction. However, we are limited in our ability to estimate the initial origins of pT181 in *S. aureus* and the timeline of transmission events due to uneven sampling and low sequence diversification. S99.5_2, a subset of CC8, constituted the first wave of MRSA in the 1960s ^14,76^, and was therefore likely sampled at high frequency relative to its true abundance within the species. While samples were collected in the pre-antibiotic era, sampling began in earnest after antibiotics were deployed clinically, with *S. aureus* samples increasing in number in the 1960s ^77^. Further conclusions about the patterns and timeline of spread taken from analysis of public genomes will need to be performed with more comprehensive sample sets, controlled for sampling bias. Samples are disproportionately taken from human hosts in hospital settings in the Global North, failing to represent the full diversity of the *S. aureus* species. Other lineages may have contained pT181 at the same time or earlier than S99.5_2, but were undetected as these strains were not frozen and later sequenced when the era of large scale “next-generation” Illumina sequencing began in the 2010s ^78^. The likely presence of pT181 in *S. aureus* before the 1950s may have reflected selection to defend against tetracyclines produced as natural products by other bacteria, such as *Streptomyces* ^1^. Environmental conditions, such as heavy-metal and pH stressors, could also have selected for pT181-containing isolates before human use of antibiotics became widespread, as TetK is able to efflux heavy metals and acts as a proton antiporter ^79,80^. There may be early collections of strains still available to be sequenced to help refine our knowledge of the genomics of this era.

By using pT181 as a model, this project uses new approaches for leveraging historical data and determining plasmid copy number. Our work raises questions around MGE evolution and lifestyles, as this single plasmid’s lifestyle has dramatically diversified since it was first detected in the 1950s and 1960s. This work adds to the knowledge base that suggests that MGEs are not as constrained to a single lifestyle, structure, and location. Going forward, our work has possible applications for others studying MGEs and antimicrobial resistance. We also show the importance of depositing sequence data from before the current era into public databases. The immediate post-WWII era was marked by a sudden, extreme chemical warfare against the most common bacterial pathogens. It is important to understand how they evolved to survive this challenge because it likely influences trade-offs in fitness in current clinical strains. Any existing unsequenced cultures of *S. aureus* collected before 1995 need to be preserved and sequenced, as they are vital historical records that can help us understand this important period in the evolution of this pathogen.

## METHODS

### pT181 genomics

*S. aureus* isolates containing pT181 were identified by BLASTN (v2.12.0+) ^47^ in the Staphopia v2 (n = 83,366) ^4^ dataset of publicly available data assembled from SRA with SKESA ^81,82^ in Bactopia (v1.7.0) ^83^. Samples were considered to contain pT181 if the sum of hit coverage exceeded 4.4 kbp (Fig. 9). Identical scripts and thresholds were used to identify pT181 in the complete genomes. Non-*S. aureus* isolates containing pT181 were identified using BLASTN ^47,84^ in the nucleotide database using the NCBI web interface (https://blast.ncbi.nlm.nih.gov/). Searches were performed on 2023-11-05 with the default parameters. We identified non-*S. aureus* sequences with >20% identity and >50% coverage with pT181. We then applied the same thresholds, as for the *S. aureus* genomes, ensuring that the sum of all pT181 coverage was >4.4 kbp. The CDS phylogeny of pT181 was created with IQ-TREE2 (v2.1.4-beta) ^85^, using a multisequence alignment generated by mapping to isolate DRX100326 with SKA (v1.0)^86^ and partitioning the CDS regions. Snp-dists (v0.8.2)^87^ was used to compare the distances calculated from a SKA (v1.0) ^86^ alignment to a reference sequence versus a reference free alignment. The calculated SNP distances were significantly different (Wilcoxon signed rank test with continuity correction, p < 2.2 x 10^-16^), though this is due in part to the large sample size for pairwise comparison. The median and modal difference between methods is 0 SNPs detected. 0.1 more SNPs were detected on average in the reference-free method than the mapping method.

Metadata of samples used in this study was downloaded from SRA for *S. aureus* isolates and from GenBank for the non-*S. aureus* isolates. Manual processing of *S. aureus* collection dates for downstream analyses was performed using RStudio (v1.3.1056) ^88^ with R (v4.0.2) ^89^ and tidyverse (v1.3.1) ^90^. We reduced the set of plasmid samples to 2,460 *S. aureus* pT181 sequences by selecting only those with a valid date, typed SCC*mec* (see “SCCmec typing”), gold rank assignment ^83^, and a genetic strain assignment ^4^.

**Fig. 9:**
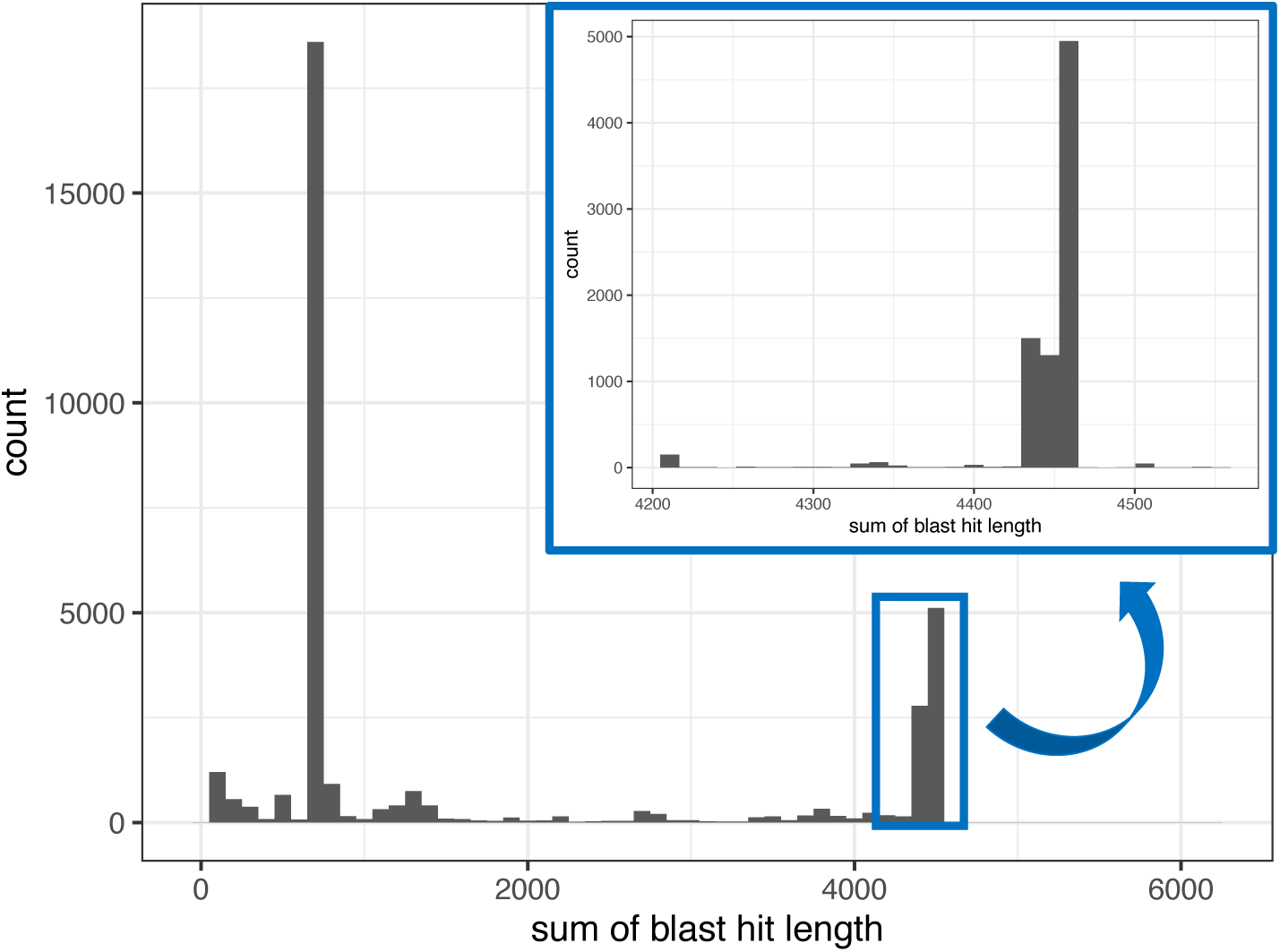
Distribution of sum of BLASTN hits to screen for pT181 presence from initial screen of subset of SRA-assembled *S. aureus* isolates (n = 82,386). Primary peak was 4425-4475 bp and the threshold for inclusion of isolates was set at 4400 bp.

### SCC*mec* typing

SCC*mec* typing was performed using sccmec (v1.2.0) ^91^. The sccmec tool compares assembled genomes to a database of known SCC*mec* elements using BLASTN. The database includes specific SCC*mec* targets (*mec*, *ccr* complex) as well as full cassettes for 15 SCC*mec* types as well as 20 subtypes. Based on the presence of SCC*mec* targets and cassette coverage, sccmec will assign a type and subtype.

### pT181 SNP calling

As pT181 is a circular replicon, it was necessary to rotate (i.e., resetting the circular plasmid to a consistent start point) the plasmid so that all of the sequences in the files would start at the same position. The start position of BLASTN (v2.12.0+) ^47^ hits to the reference plasmid were used to identify an identical start position. Samtools (v1.9 with htslib v1.9) ^92^ faidx was used to extract these positions. Picard (v3.1.1) ^93^ normalize was used to generate fasta files with equal line lengths. SNPs, small indels, and complex mutations were identified using Snippy (v4.6.0) ^48^. Mutation spikes were not due to our rotation of the plasmid for alignment. The mutations that occur in the spike positions occur regardless of where the start of the contig is located (Fig. S4). Gene positions were identified with bakta (v1.8.2) ^94^ annotation and gene plot (Fig. 2) was visualized with gggenes^95^.

### Testing for recombination

Linkage analysis was performed via a PHI test ^54^ run with PhiPack (v1.1) ^96^, on a reference-free SKA (v1.0) ^86^ alignment of all 7,988 pT181 sequences.

### Plasmid copy number

**Fig. 10:**
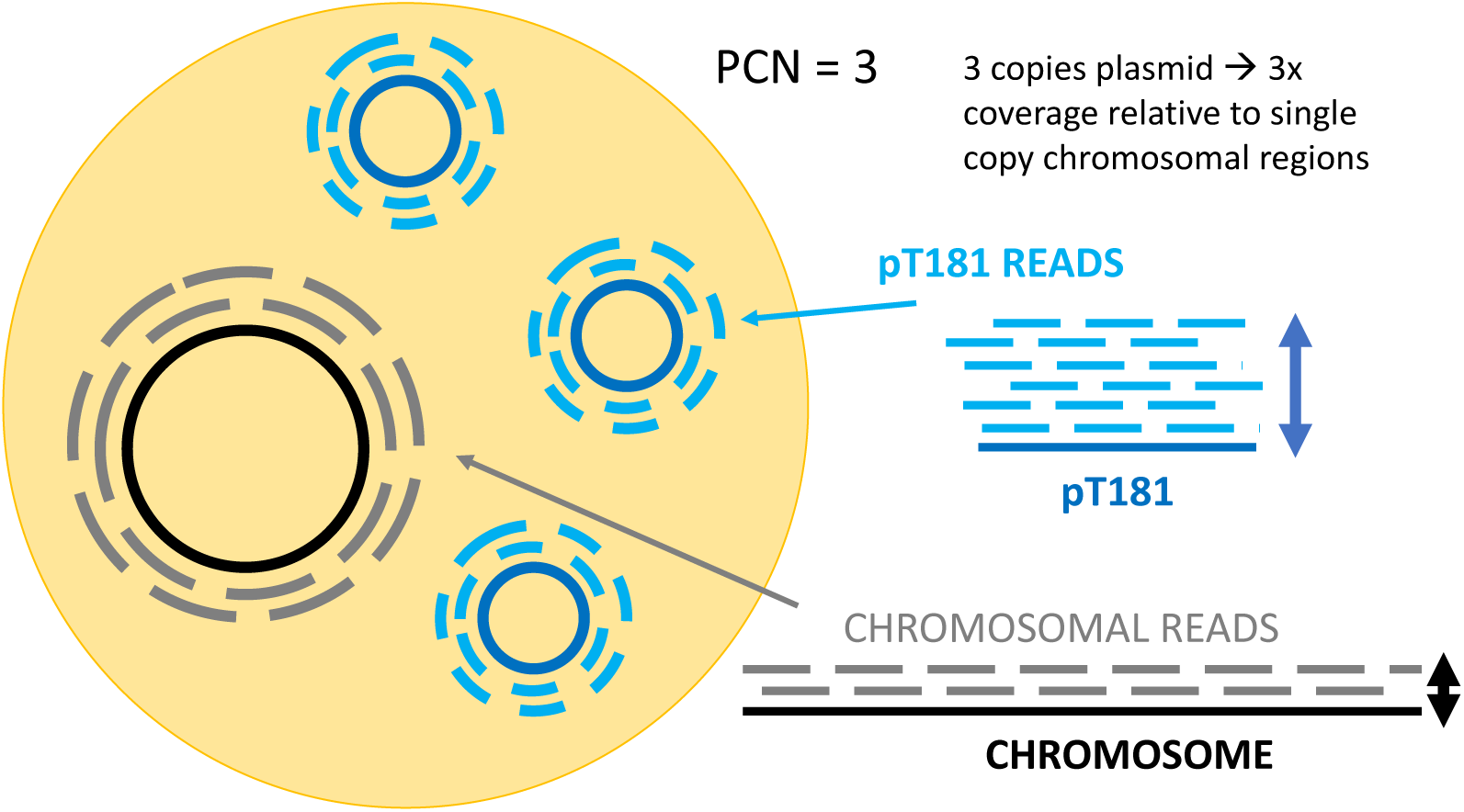
Cartoon of read mapping and relative read depth measurements. The relative read depth of the plasmid provides an estimate of its copy number.

Illumina reads from each isolate were mapped with BWA (v0.7.17) ^97^ to each isolate’s own assembled genome (Fig. 10). Read depth across the genome was calculated with samtools depth at all sites (v1.9, using htslib v1.9) ^92^. Single-copy chromosomal contigs were identified by the presence of known single copy chromosomal genes, found with BLASTN (v2.12.0+)^47^. To ensure an accurate ratio, only assemblies ranked as “gold” by Bactopia ^83^ were used for the PCN analysis.

PIRATE (v.1.0.5) ^98^ gene families were used to identify single copy chromosomal genes. We utilized gene families found in >95% of genomes at exactly 1 copy per genome, with fewer than five fission loci.

The cutoff for single-copy pT181 (PCN < 2) was validated using the distribution of coverage for contigs containing single-copy core genes for gold-ranked isolates (Fig 11). The distribution of chromosomal copy number (CCN) ranges was centered on 1, but values range from less than 0.5 to greater than 2. To determine the copy number of genes located near the origin, including SCC*mec*, we calculated average read depth of contigs containing one of the four genes (*rpmH, dnaA, dnaN,* and *recF*) located near the origin of replication ^99^. The distribution of CCN for those genes was slightly higher (mean = 1.15, sd = 0.11). This distribution had a similar center to the distribution of pT181 for SCC*mec* type III, which has the pT181 plasmid integrated into the cassette (mean = 1.16, sd = 0.057).

**Fig. 11:**
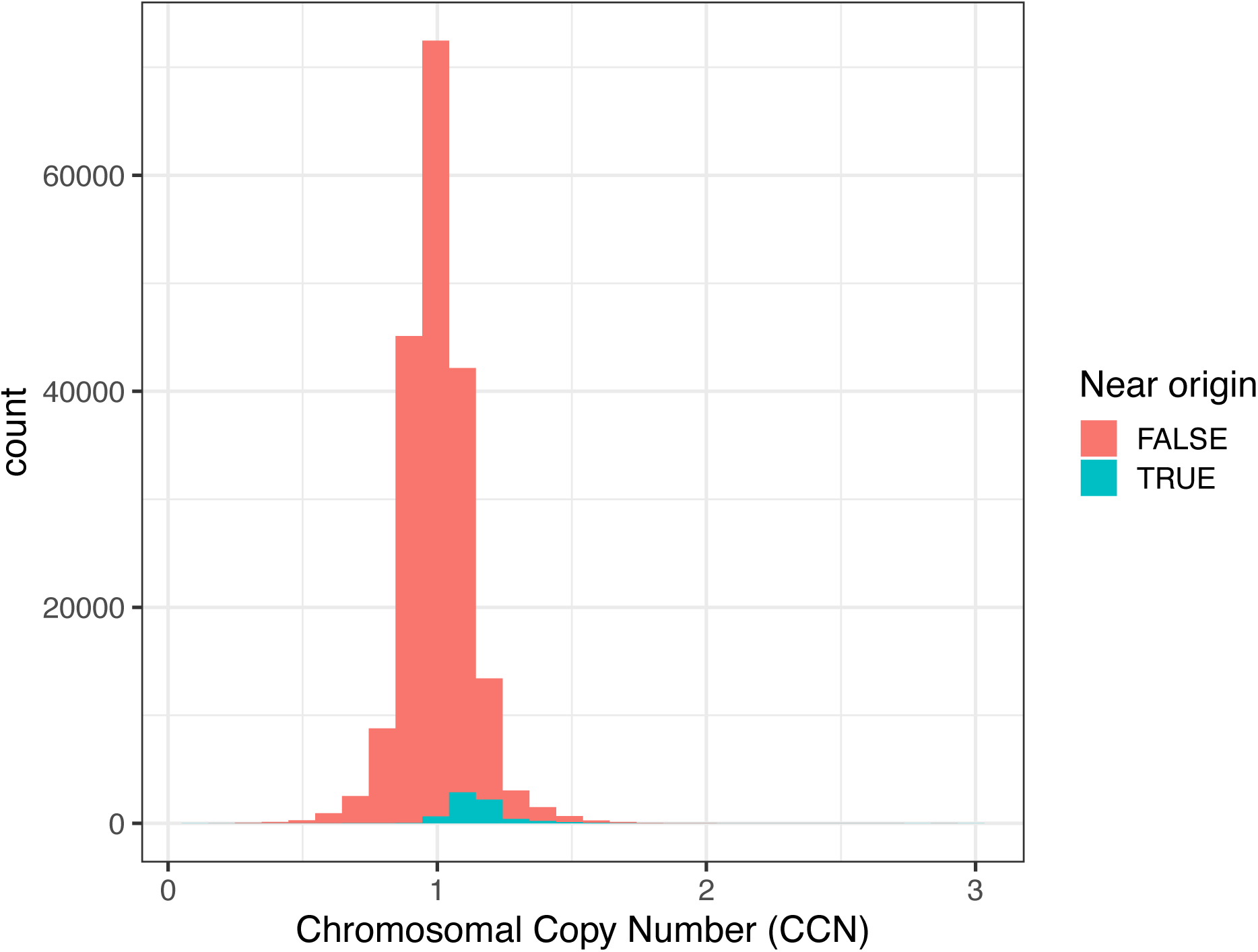
Distribution of chromosomal copy number (CCN), colored by position relative to origin of replication. Contigs are considered near the origin if they contain at least one of the 4 genes (*rpmH, dnaA, dnaN,* and *recF*) known to be close to the origin of replication^99^

We assessed the coefficient of variation, or the standard deviation of read depth divided by mean read depth, for each contig for a test set of 10 isolates (Fig. 12). Core chromosomal and pT181 contigs are not significantly different in their coefficient of variation (Wilcoxon rank sum, p = 0.28). This analysis convinced us that read mapping variation across pT181 was not significantly different from chromosomal read mapping, allowing us to use this process for copy number analysis.

**Fig. 12:**
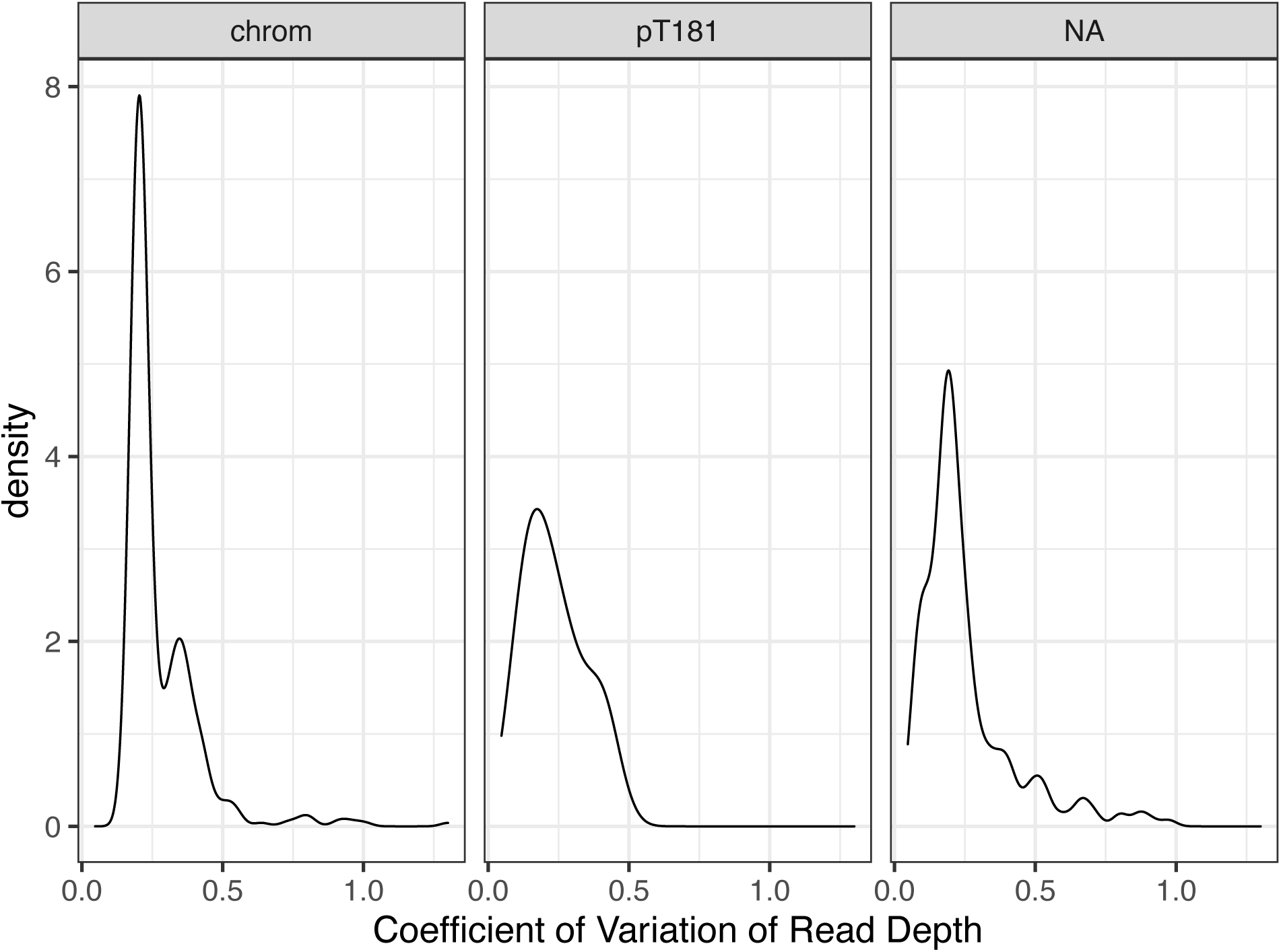
Coefficients of variation of read depth (standard deviation divided by mean) for known chromosomal contigs, pT181, and other contigs follow a similar distribution across 10 test gold-quality isolates.

### PCN as a function of time and strain

PCN over time was assessed using a Gamma (log link) generalized linear mixed effect model (GLMM) with R (v4.4.2) ^89^ using the package glmmTMB (v1.1.11) ^100^. Strains and BioProjects containing at least 5 multicopy isolates were selected; to be used, a sample must belong to a strain and BioProject combination containing at least 4 other samples (n = 455). Post hoc testing was performed with the emmeans package (v1 11.1) ^101^.

Year and strain were assessed as fixed effects, as we are interested in both of their impacts on PCN. BioProject was assigned as a random effect within the fixed effects of strain and year. This is to account for possible technical variation between projects, while recognizing that BioProjects frequently contain one or few strains and are typically collected in a narrow time frame. Formula for glmmTMB call is as follows: PCN ∼ numYear + strainFac + (1 | strainFac:BioProject) + (1 | numYear:BioProject). numYear refers to year of collection as a continuous variable and strainFac refers to strain as a factor.

**Fig. 13:**
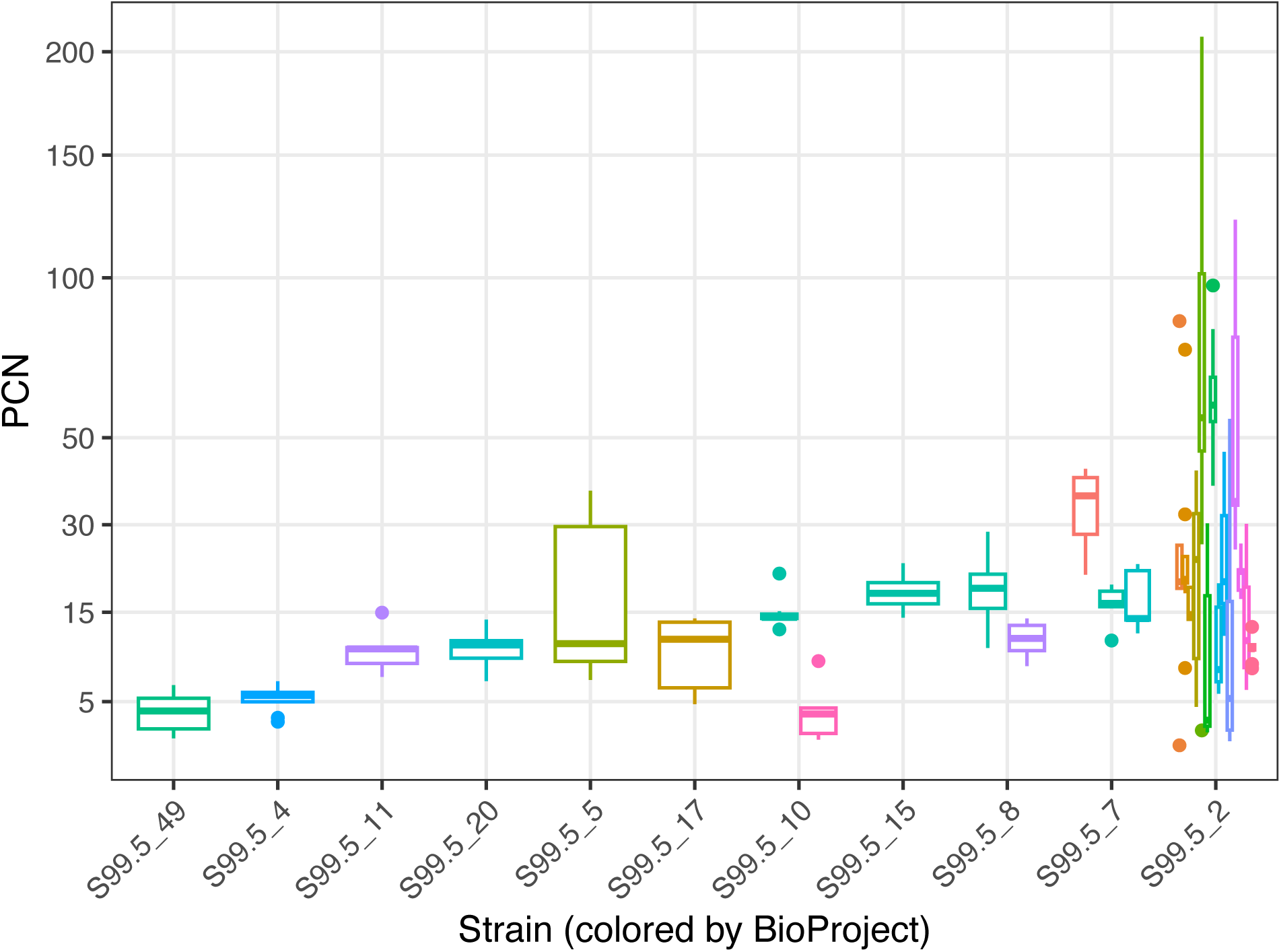
Strains vary in median copy number, with an effect of BioProject. Each BioProject has a different boxplot within the strain. Data shown for strain x BioProject combinations containing at least 5 isolates.

### Investigating impact of extraction technique

Technical factors, such as extraction technique, have been previously documented to impact PCN estimates^102^. Given that this is a retrospective genomic epidemiology paper, we are not able to control for the conditions these samples were collected and sequenced under. We performed a literature search to identify publications associated with BioProjects containing samples collected before 1995 or containing at least 10 multicopy isolates. We identified 9 BioProjects with usable extraction techniques and categorized them into 3 categories: silica, salt, or membrane of unspecified chemistry. All early (pre-1995) samples for which we have extraction technique are S99.5_2 isolates extracted with silica-based protocols and sequenced on an Illumina HiSeq 2000 at Wellcome Sanger Institute; we compared the PCN of these samples to post-1995 samples of the same strain, extraction technique, instrument, and center. All of these samples, collected pre- and post-1995 (n = 180), were released 2012-2013. This does not tell us for certain when they were processed, but it was likely close enough in range that we would expect reasonable consistency in techniques within a sequencing center over that time period. There has been a marked decline in multicopy PCN over the collection period (1960 - 2010) for samples with these identical conditions (linear regression, p = 5.9 x 10^-16^). Moreover, it does not appear that the extraction technique has a significant impact on our PCN estimates. If we exclude older samples, which are all silica-based, and focus on isolates post-1995 for which we have extraction technique, extraction technique does not have a significant relationship with copy number (linear regression, salt-based as reference; p = 0.10 silica-based; p = 0.26 membrane of unspecified chemistry).

### Identifying pT181 novel chromosome insertion sites

Using the core gene BLASTN hits used to identify known single copy genes, as well as searches for pT181, we looked for contigs that contained both pT181 and core genes. 19/7,927 *S. aureus* pT181 SRA isolates were found to have core genes on the same contig as pT181. We used Prodigal ^103^ annotations generated with Bactopia (v1.7.0) ^83^ to identify the genes directly surrounding the insertion site of the plasmid. Genes from these hits, as well as a complete genome, were visualized using clinker (v0.0.31) ^104^.

The majority of pT181-containing contigs were the same canonical length of the plasmid, approximately 4,440 kbp. We are likely unable to identify genes on other contigs of most putatively integrated pT181, due to repetitive sequences making it difficult to assemble in those regions.

### Binning samples by time

There were no samples in the n = 2,460 set collected 1992-1996. This lull in high-quality samples formed a natural distinction between these two groups, given the difference in sample composition and plasmid frequency in pre- and post-1995 pT181 isolates.

## Data availability

All genomes used in this study are publicly available. The 61 non-*S. aureus* accessions are included as a supplementary file. The summary file containing *S. aureus* SRA accessions is available on Zenodo at https://zenodo.org/records/10471309 (staphopia_v2_all_report.txt). The 337 complete *S. aureus* genome accessions with PIRATE gene annotations are available at https://github.com/VishnuRaghuram94/SCAPE/blob/main/timsfigsData.zip (dnaA_start_coords.gfs). Scripts used in analysis and creating figures are available at https://github.com/phillma2/pT181.

## Acknowledgements

M.A.P. was supported by funding from NSF 2146260, NIH awards AI158452, AI153116, and T32 AI138952 (Infectious Disease Across Scales Training Program). R.A.P. III & T.D.R. were supported by the Office of Advanced Molecular Detection, Centers for Disease Control and Prevention cooperative agreement number CK22-2204 through contract 40500-050-23234506 from the Georgia Department of Public Health. D.B.W. was supported by funding from the Sloan Foundation (Sloan Research Fellowship FG-2021-16667) and NSF 2146260. T.D.R. was supported by funding from NIH awards AI158452 and AI153116. We thank Vishnu Raghuram and Ananya Saha for discussions about the manuscript.

## Supplementary figures

**Fig. S1:**
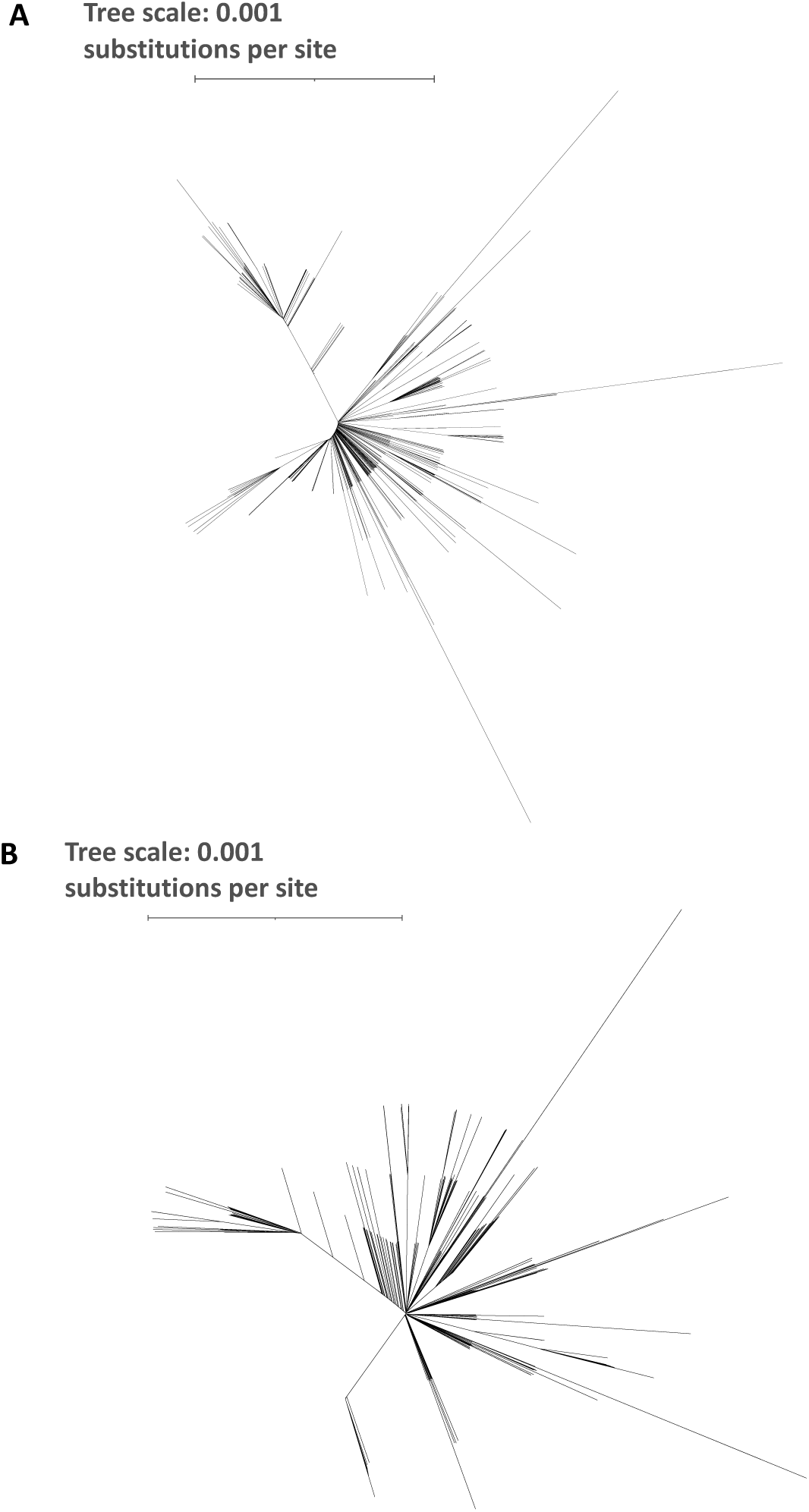
Unrooted phylogenetic trees for pT181 A) whole plasmid sequence and B) CDS. Scale is in substitutions per site.

**Fig. S2:**
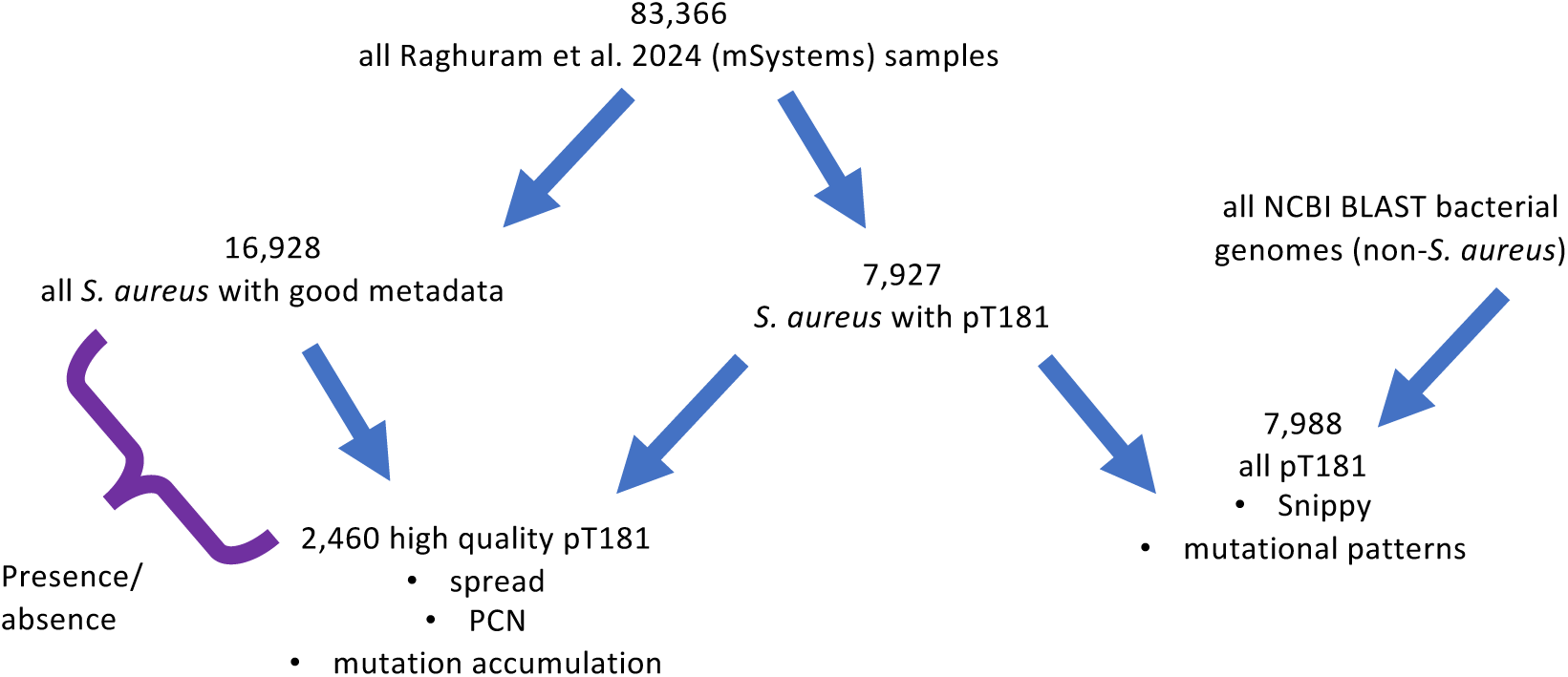
Diagram of sample filtering steps

**Fig. S3:**
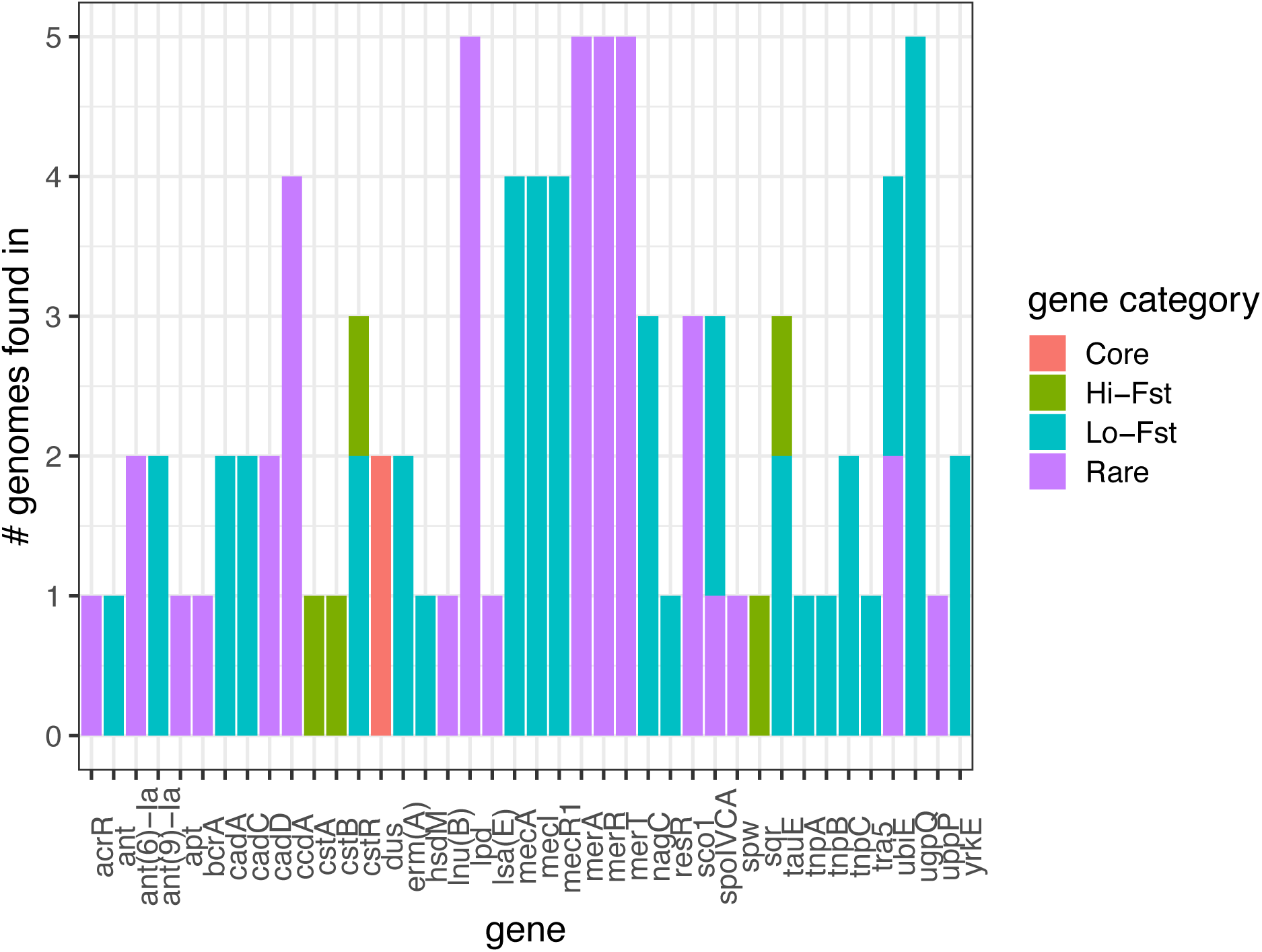
pT181 integrates into a region of the chromosome consisting primarily of strain-diffuse and rare genes ^4^. No genes present in positions 55-90kbp in complete genomes with chromosomally integrated pT181 are universal to this region in all isolates.

**Fig. S4:**
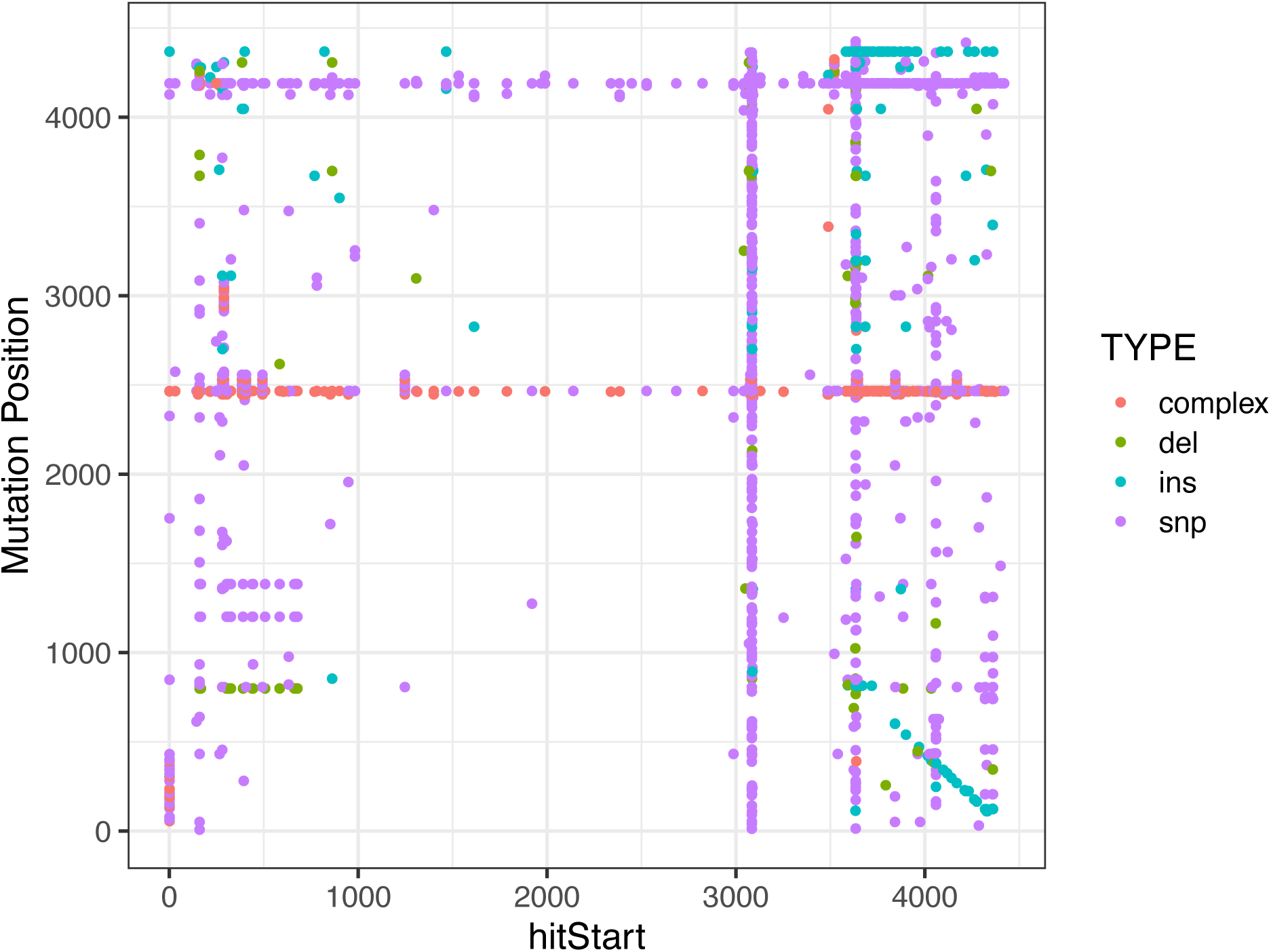
Pre-rotation start position of plasmid contig does not influence position of mutations detected in pT181. X-axis, hitStart, indicates start position of isolates’ genomes relative to blast query (i.e., subject start / “sstart” for retrieval with blastn). Y-axis, mutation position, indicates where in the sequence mutations occurred, following their rotation to have equal start points. The presence of the mutational spikes across a range of hitStart positions indicates that it is not the rotation positions that are generating these spikes; they are biological in origin and not due to technical error.

